# Cancer-specific associations of driver genes with immunotherapy outcome

**DOI:** 10.1101/2020.06.16.155895

**Authors:** Tomi Jun, Tao Qing, Guanlan Dong, Maxim Signaevski, Julia F Hopkins, Garrett M Frampton, Lee A Albacker, Carlos Cordon-Cardo, Robert Samstein, Lajos Pusztai, Kuan-lin Huang

## Abstract

Genomic features such as microsatellite instability (MSI) and tumor mutation burden (TMB) are predictive of immune checkpoint inhibitor (ICI) response. However, they do not account for the functional effects of specific driver gene mutations, which may alter the immune microenvironment and influence immunotherapy outcomes. By analyzing a multi-cancer cohort of 1,525 ICI-treated patients, we identified 12 driver genes in 6 cancer types associated with treatment outcomes, including genes involved in oncogenic signaling pathways (NOTCH, WNT, FGFR) and chromatin remodeling. Mutations of *PIK3CA, PBRM1, SMARCA4*, and *KMT2D* were associated with worse outcomes across multiple cancer types. In comparison, genes showing cancer-specific associations—such as *KEAP1, BRAF, and RNF43*—harbored distinct variant types and variants, some of which were individually associated with outcomes. In colorectal cancer, a common *RNF43* indel was a putative neoantigen associated with higher immune infiltration and favorable ICI outcomes. Finally, we showed that selected mutations were associated with PD-L1 status and could further stratify patient outcomes beyond MSI or TMB, highlighting their potential as biomarkers for immunotherapy.

## Introduction

Immune checkpoint inhibitors (ICIs) have dramatically transformed the treatment landscape of advanced cancers. For some patients, these agents extend life and provide durable benefits. However, the majority of patients do not benefit from ICIs; response rates in clinical trials range from 10-50%, and a significant fraction of patients suffer from immune-related adverse events^1,2^. There is an urgent need for reliable predictors to better select patients for ICI therapy.

Several ICI biomarkers have been identified, but only a few are in routine clinical use: PD-L1 expression levels in tumor and immune cells, and deficient mismatch repair (dMMR)/ microsatellite instability-high (MSI-H) have been incorporated into FDA-approved indications for some ICIs. However, these are still imperfect predictors; response rates among MSI-H patients in the clinical trials leading to the FDA approval of this biomarker were just under 40%^3^. Tumor mutation burden (TMB) is emerging as another predictive biomarker but has not been widely implemented due to a lack of standardized definitions and thresholds^4^. Retrospective studies have suggested that combinations of TMB, MSI, and PD-L1 can be used to further stratify patients^5–8^. Biomarkers based on gene expression profiles or other aspects of the tumor microenvironment have also been proposed^9,10^.

Driver gene mutations are reliably measured across sequencing panels and observed in significant fractions of cancer cases. Cancer driver genes alter tumor cell phenotypes in ways that may modulate tumor-immune interactions. Mutations in *STK11*^11^, *KEAP1*^12,13^, *JAK1, JAK2,* and *B2M*^14^ have been associated with ICI resistance, while mutations in *PBRM1*^15,16^, *POLE*, *POLD1*^17^, *SERPINB3*, and *SERPINB4*^18^ have been associated with ICI benefit. However, previous studies seeking mutations predictive of ICI benefit have been limited to relatively small cohorts in a few cancer types^15,19,20^, and have rarely explored predictive factors beyond TMB/MSI^5–7,21^, nor have they examined the predictive role of gene mutations at the variant level^11,12^.

In this study, we applied statistical modeling to a multi-cancer dataset of 1,525 ICI-treated patients to identify mutated driver genes associated with cancer-specific ICI benefit or resistance. We further analyzed these immunotherapy-response associated genes (IRAGs) at the variant level, and correlated them with features of the tumor immune microenvironment. Based on these analyses, we observed that the IRAGs improved prediction of ICI outcomes over MSI or TMB alone, which suggests they may have immediate clinical relevance.

## Results

### Clinical and genomic characteristics of the multi-cancer cohort

For discovery, we utilized the multi-cancer cohort from Samstein et al. (Figure 1A)^21^. After excluding cancer types with fewer than 20 cases, the discovery cohort consisted of 1,525 patients who had next-generation sequencing profiling of their tumors and received at least one dose of an ICI. The three most common cancer types in the discovery cohort were melanoma (SKCM, n = 300), lung adenocarcinoma (LUAD, n = 297), and bladder cancer (BLCA, n = 215)(**Supplemental Table 1**). The cohort was predominantly male (63%), with a quarter of patients over the age of 70; the majority of patients were treated with an anti-PD-1/PD-L1 agent (79%), while the rest received either anti-CTLA-4 therapy (6%), or a combination (15%). For validation, we two independent cohorts, Miao et al.^20^ and Van Allen et al.^19^, described further in the Methods.

**Figure 1.**
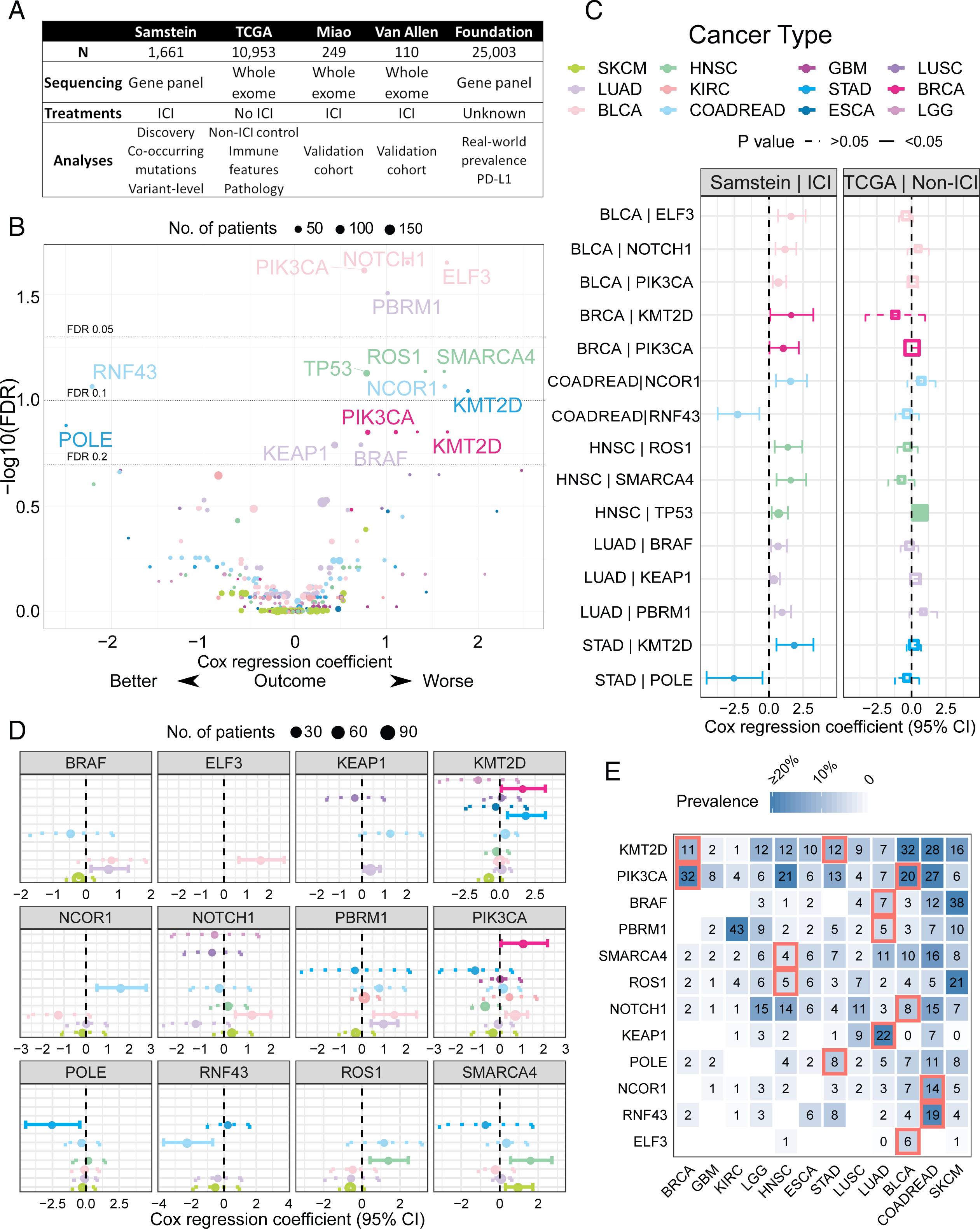
Clinical and genomic characteristics of the cohort. (A) Cohorts used in this study, indicating the number of patients, sequencing technology used, immune checkpoint inhibitor (ICI) exposure, and the analyses for which each cohort was used. (B) Immunotherapy-response associated genes (IRAGs) were identified using multivariable Cox proportional hazards models stratified by cancer type and corrected for age, sex, ICI class, and TMB. This volcano plot shows the negative log_10_ false discovery rate (FDR) plotted against regression coefficients for each mutated driver gene. Labelled genes are IRAGs with p<0.05 and FDR<0.2. Negative coefficients indicate better overall survival among mutated tumors. The same colors and abbreviations are used to refer to cancer types throughout the manuscript. BLCA: Bladder carcinoma; BRCA: Breast carcinoma; COADREAD: Colorectal adenocarcinoma; ESCA: Esophageal carcinoma; GBM: Glioblastoma; HNSC: Head and neck squamous cell carcinoma; KIRC: Renal cell carcinoma; LUAD: Lung adenocarcinoma; LUSC: Lung squamous cell carcinoma; LGG: Low-grade glioma; STAD: Stomach adenocarcinoma; SKCM: Melanoma. (C) Comparison of Multivariable Cox regression coefficients for IRAGs identified in the ICI-treated Samstein cohort as opposed to the non-ICI TCGA cohort (models stratified by cancer and corrected for age, sex and TMB). IRAGs associated with survival only in ICI-treated patients were considered predictive rather than prognostic. Solid lines and filled shapes indicate p<0.05, while dashed lines and open shapes indicate p>0.05. The size of points reflects the number of patients with the mutation. (D) Cox proportional hazards regression coefficients with 95% confidence intervals of IRAGs across all cancer types in the Samstein cohort. Suggestive associations (p<0.05) are shown with solid lines and filled shapes; dotted lines and open shapes indicate p>0.05. The size of points reflects the number of patients with the mutation. Negative coefficients indicate improved overall survival among mutated tumors. (E) The prevalence of each IRAG across cancer types in the Samstein cohort is represented colorimetrically; darker blue indicates a higher mutation prevalence. Red outlines indicate the cancer types in which the IRAGsweresignificantlyassociatedwithoverallsurvival(p<0.05, FDR<0.2).

### Genes associated with immunotherapy outcomes

To identify associations between individual mutated genes and overall survival, we constructed multivariable Cox regression models for each cancer type, adjusting for patient age, sex, ICI class and TMB (**Methods**). After correcting for multiple hypothesis testing, we identified 13 genes in 6 cancer types associated with overall survival (FDR< 0.2), we call these genes ICI response/resistance-associated genes (IRAGs) (Figure 1B and **Supplemental Table 2**). These IRAGs comprised genes involved in chromatin remodeling (*NCOR1, SMARCA4, KMT2D, PBRM1*), oncogenic signaling pathways (*PIK3CA, BRAF*), NOTCH (*NCOR1, NOTCH1, PBRM1*) and WNT (*RNF43, SMARCA4, KMT2D*) pathways, and *TP53*.

To rule out the possibility that mutations in these genes are prognostic rather than predictive of ICI benefit, we repeated the multivariable regression analysis in the TCGA cohort, which does not include ICI-treated patients (Figure 1A, 1C)^23^. Since *TP53* mutations were significantly associated with worse outcomes regardless of treatment received, they were considered likely prognostic rather than predictive, and were excluded from further evaluation.

In order to validate the IRAGs, we examined their associations with clinical benefit in two independent ICI-treated cohorts: 249 microsatellite stable cases published by Miao et al.^20^ and 110 SKCM cases published by Van Allen et al.^19^ (Figure 1A). Bearing in mind the clinical differences between these cohorts, a few IRAGs exhibited consistent associations across the studies: *RNF43, ROS1, POLE, and KEAP1*(**Supplemental Figure 2**). More ICI-treated cohorts representing a greater variety of cancer types are needed to thoroughly validate the IRAGs.

### Cancer-specificity of immunotherapy response-associated genes

Next, we asked whether these IRAGs showed similar predictive value across different cancer types. Four IRAGs had were associated (p<0.05) with overall survival in more than one cancer type: *KMT2D* in breast (BRCA, HR = 5.3, p = 0.04) and stomach cancers (STAD, HR = 6.7, p = 0.005); *PBRM1* in BLCA (HR = 4.5, p = 0.002) and LUAD (HR = 2.8, HR, p<0.001); *PIK3CA* in BLCA (HR = 2.2, p = 0.002) and BRCA (HR = 3.0, p = 0.04); *SMARCA4* in head and neck cancer (HNSC, HR = 5.1, p = 0.002) and SKCM (HR = 2.7, p = 0.005) (Figure 1D). These associations were consistently deleterious (HR≥1). The remaining IRAGs only exhibited significant associations with survival in one cancer type in the cohort.

These cancer-type specific associations may be explained by 1) uniform effects across cancer types, but a lack of power to detect associations in cancers with low IRAG mutation prevalence or limited sample sizes, or 2) cancer-specific functional effects of IRAGs. In the first scenario, we would expect to find IRAGs significantly associated with ICI outcomes in the cancer types in which they are most commonly mutated, since these cohorts have the greatest statistical power to detect a potential association. This was the case for *ELF3* (mutated in 6% of BLCA), *RNF43* and *NCOR1* (mutated in 19% and 14% of COADREAD, respectively), and *KEAP1* (mutated in 22% of LUAD) (Figure 1E). However, the other IRAGs were not consistently associated with ICI-benefit/resistance in cancer types with the highest mutation prevalence. For example, *PBRM1* was not associated with survival in KIRC (mutated in 43%; HR = 1.2, p = 0.57), but was associated with worse survival in both LUAD (mutated in 5%; HR = 2.8, p<0.001) and BLCA (mutated in 3%; HR = 4.5, p = 0.002). These results suggest cancer-specific effects for these IRAG mutations.

### Variant-level analysis of genes associated with immunotherapy benefit

Given the cancer-specificity of certain IRAGs (Figure 1D), we hypothesized that the ICI-benefit/resistance phenotype might be a result of specific variants or classes of variants (e.g. frameshift, missense, etc.) that were unevenly distributed across the different cancer types. Accordingly, we found that the variant classes of *ELF3* in BLCA, *RNF43* in COADREAD and *KMT2D* in STAD were significantly different from those in other cancers (Chi-squared FDR = 0.03, 0.03, 0.03, respectively); for example, *ELF3* frameshift mutations were only observed in BLCA, and *RNF43* frameshift mutations were more common in COADREAD than in other cancers (Figure 2A, **Supplemental Table 3**).

**Figure 2.**
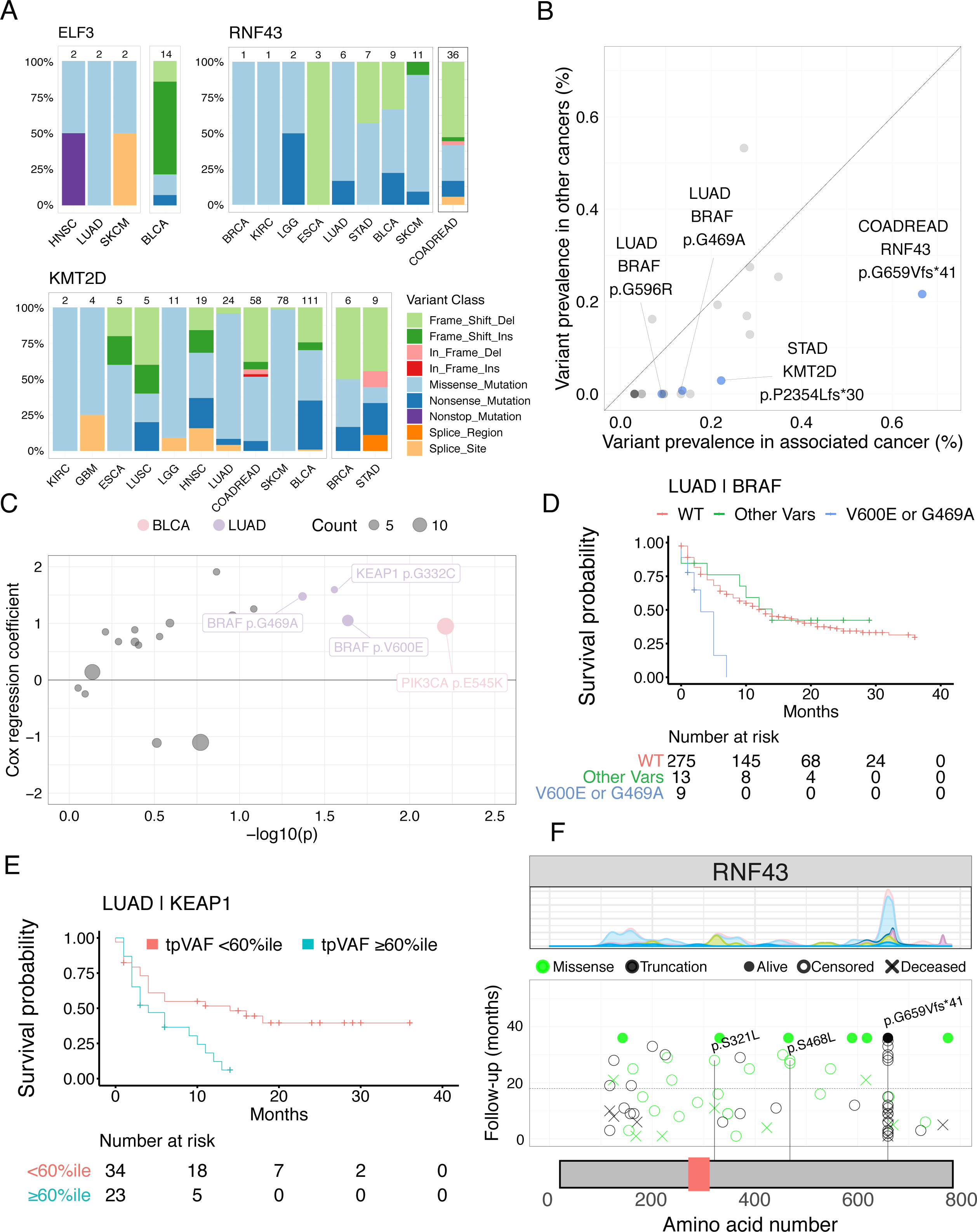
Immunotherapy-response associated variants. (A) The distribution of variant classes for *ELF3* in *BLCA*, *RNF43* in COADREAD and *KMT2D* in STAD were significantly different compared to other cancer types (Chi-squared FDR = 0.03,0.03, 0.03, respectively). (B) Scatter plot comparing the prevalence of IRAG variants in their corresponding cancer type versus other cancers with the IRAG mutation. Labelled variants were significantly more common in the IRAG cancer type than other cancers (one-sided Fisher’s exact test p<0.05 and FDR<0.2). Significant variants are colored blue; other variants are colored grey. Only variants with more than 1 occurrence in the IRAG-associated cancer were considered. (C) The variant-level analysis revealed individual IRAG variants that were independently associated with survival. This plot shows the Cox proportional hazards regression coefficients versus the level of significance (-log_10_p) for individual IRAG variants. Models were stratified by cancer type and corrected for age, sex, ICI type, and TMB. Variants with p<0.05 are labelled and colored according to the cancer type. Point size reflects the number of patients with the variant. Negative coefficients indicate better overall survival associated with the presence of the variant. (D) Overall survival was worse among LUAD patients with either BRAF variant V600E or G469A, compared to those with other *BRAF* variants (Other Vars, log-rank p = 0.005), and those without BRAF variants (WT, log-rank p<0.001). (E) Overall survival was worse among KEAP1-mutated LUAD patients with a tumor purity-adjusted variant allele fraction (tpVAF) above the 60^th^ percentile (log-rank p-value = 0.008). (F) Surviplot (survival by variant) of RNF43. The top panel is a stacked density plot showing the distribution of variants across the RNF43 protein in the entire cohort; colors represent different cancer types. The middle panel plots individual patients’ duration of follow-up (y-axis) and survival outcomes (shapes) against the amino acid position (x-axis) of each patient’s *RNF43* variant. Variants with more than one occurrence are labelled and marked with a vertical line. He dotted horizontal line is the median overall survival of all *RNF43* WT patients. The bottom panel is a schematic of the *RNF43* protein. The red box represents a RING-type zinc finger domain, as annotated by UniProt.

We predicted that different IRAG variants would have different cis-expression consequences, and explored this using TCGA gene expression data (**Methods**). Missense variants in *KEAP1* LUAD were associated with increased gene expression (FDR<0.001), while truncating variants in *KEAP1* LUAD, *PBRM1* LUAD, *RNF43* COAD, and *NCOR1* COAD were all associated with decreased gene expression (FDR = 0.06, 0.06, 0.06, 0.18, respectively) (**Supplemental Table 4**).

Next, we explored whether particular IRAG variants were more likely to occur in their IRAG-specific cancer type than other cancer types. We hypothesized that such “enriched” variants might be associated with the ICI-benefit/resistance phenotype. Using the one-sided Fisher’s exact test, we identified four IRAG variants significantly or suggestively enriched in their corresponding cancer type: *RNF43* p.G659Vfs*41 in COADREAD (OR = 7.0, FDR = 0.02); *BRAF* p.G469A (OR = 21.0, FDR = 0.10) and p.G596R (OR = Inf, FDR = 0.15) in LUAD; and *KMT2D* p.P2354Lfs*30 (OR = 9.3, FDR = 0.20) in STAD (Figure 2B, **Supplemental Table 5**). Interestingly, the *RNF43* p.G659Vfs*41 variant has been identified as an immunogenic neoantigen and a possible target for cancer vaccines.^24,25^ Using PepQuery^26^, we confirmed that thep.G659Vfs*41 variant was expressed at the peptide level in mass-spectrometry proteomic data of COADREAD tumor samples (**Supplemental Figure 6, Supplemental Table 17**).

To further test the hypothesis that specific variants may be associated with specific ICI-benefit/resistance phenotypes, we conducted multivariable Cox regression analysis at the level of individual variants within the IRAGs, stratified by cancer type (**Methods**). Although power was limited, *PIK3CA* p.E545K in BLCA was significantly associated with worse ICI outcomes (HR = 2.6, FDR = 0.02). Several suggestive associations (p<0.05) were also identified: *BRAF* p.V600E (HR = 2.9, p = 0.02); *BRAF* p.G469A (HR = 4.4, p = 0.04); and *KEAP1* p.G332C (HR = 4.9, p = 0.03) in LUAD (Figure 2C, **Supplemental Table 6**). LUAD samples with either *BRAF* p.G469A or p.V600E hotspot mutations exhibited worse outcomes compared to other *BRAF*-mutated LUAD (log-rank p = 0.005) and wild-type (WT) LUAD (log-rank p<0.001) (Figure 2D). The *RNF43* p.G659Vfs*41 variant showed a tendency towards improved overall survival, though the association was not significant (HR=0.33, p=0.17)(Figure 2F, **Supplemental Table 6**).

We hypothesized that the clonality of an IRAG mutation might further influence ICI outcomes. Since there were very few copy number alterations involving IRAGs (6 in total), we reasoned that tumor-purity-adjusted variant allele fraction (tpVAF) could be a proxy for mutation clonality. Although continuous tpVAF was not associated with overall survival in multivariable Cox regression models, a binary cutoff of *KEAP1* tpVAF ≥60^th^ percentile (optimized using receiver operating characteristic analysis, **Supplemental Table 7**) was independently associated with worse overall survival in LUAD (HR = 2.2, FDR = 0.18)(Figure 2E, **Supplemental Table 8**).

### Dissecting the effects of co-occurring mutations

Specific combinations of driver gene mutations co-occur in cancer more frequently than expected by chance^27^, potentially providing a synergistic selective advantage or arising from a shared mutagenic process. Mutually exclusive mutations have also been described ^28,29^, often due to redundant functional impacts on shared pathways. Co-occurring or mutually exclusive mutations may confound or interact with the associations we identified between IRAGs and outcomes after immunotherapy. To explore this possibility, we used two-tailed Fisher’s exact tests to identify genes with a significant tendency to co-occur or be mutually exclusive with each IRAG in their associated cancer types (**Methods**). Six IRAGs had significantly co-occurring mutations (FDR<0.05): *KEAP1* and *PBRM1* in LUAD, *NOTCH1* in BLCA, *KMT2D* in STAD, and *NCOR1* and *RNF43* in COADREAD (**Supplemental Table 9**). Of note, the IRAGs *NCOR1* and *RNF43* co-occurred with each other in COADREAD (OR = 4.9, FDR = 0.05). The only significant mutual exclusivity was observed between *RNF43* and *TP53* in COADREAD (OR=0.11,FDR=0.009).

Next, we examined whether IRAG associations were independent of the co-occurring mutations. *KEAP1* in LUAD remained an independent predictor of survival even after accounting for *STK11* (HR = 1.5, p = 0.05). *RNF43* (HR = 0.08, p = 0.001) and *NCOR1* (HR = 9.0, p<0.001) in COADREAD were also both independently associated with survival, even after accounting for the presence of the other mutation (**Supplemental Table 10**).

We also tested for interactions between co-occurring mutations, and found a significant interaction between *STK11* and *KEAP1* (HR = 2.4, p = 0.03 for the interaction, **Supplemental Table 10**). Median overall survival was significantly worse among patients with *KEAP1*+*STK11* tumors compared to *KEAP1* alone (4 vs. 14 months, log-rank FDR = 0.03), *STK11* alone (4 vs. 18 months, log-rank FDR = 0.03), or WT (4 vs. 13 months, log-rank FDR = 0.01) (Figure 3F). This finding was replicated in the Miao cohort, where we found that *KEAP1*+*STK11* tumors had significantly worse outcomes than WT tumors (HR = 66.1, FDR = 0.02) (**Supplemental Table 11**).

**Figure 3.**
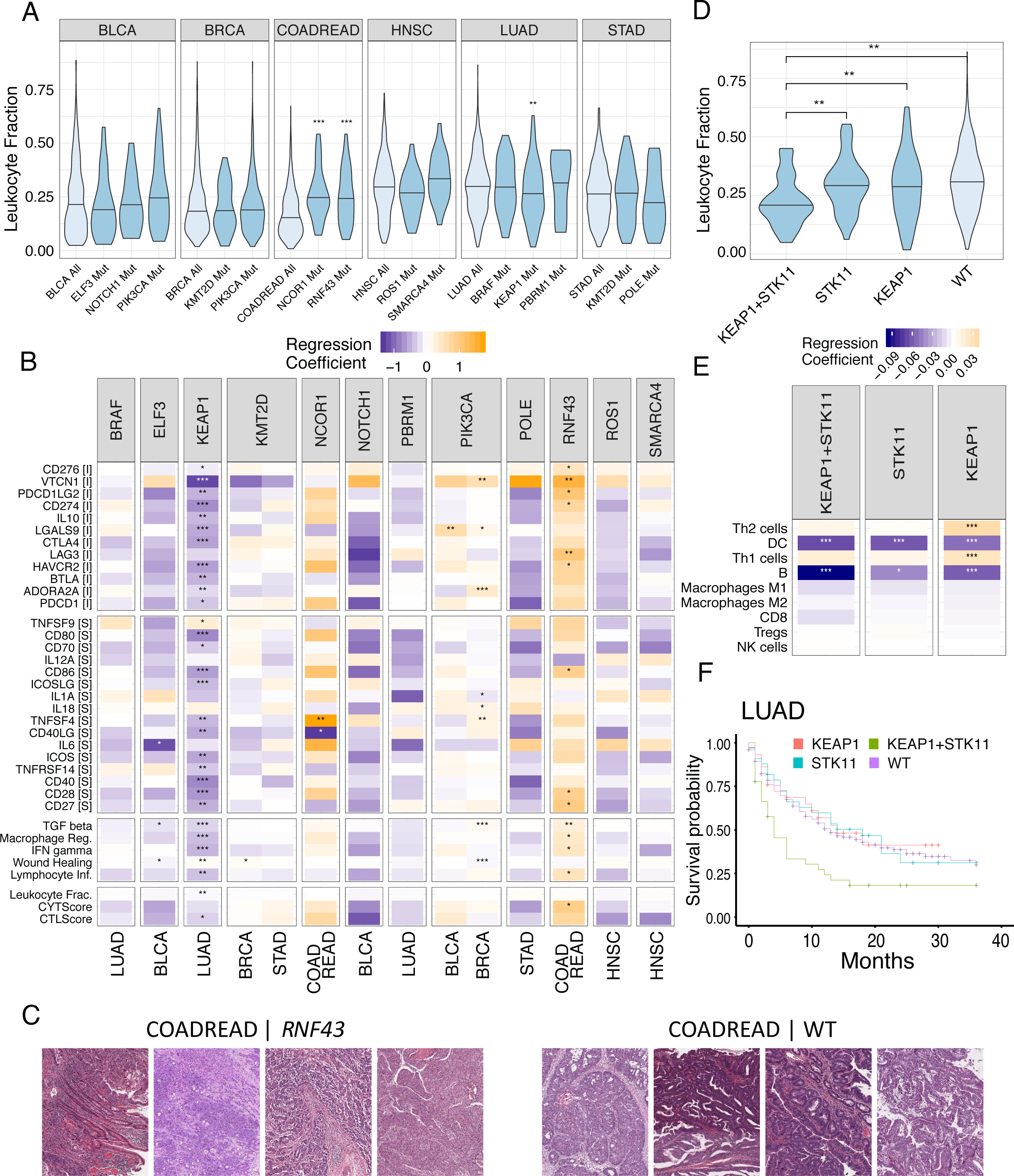
Immune microenvironment associated with IRAG mutations. (A) Violin plots showing the leukocyte fraction of IRAG-mutated tumors in TCGA. The horizontal line indicates the median for each population. Comparisons were made using the Wilcoxon rank-sum test. Significance levels of the associations are labelled as FDR<0.15 (*), FDR<0.05(**), FDR<0.01 (***). (B) Color gradients representing linear regression coefficients for the associations between IRAG mutation status and immune features in TCGA, where white represents no change, orange represents IRAG mutations associated with up-regulation, and blue represents IRAG mutations associated with down-regulation. Significance levels of the associations are labelled as FDR<0.15 (*), FDR<0.05(**), FDR<0.01 (***). Immunomodulators are labelled [I] for inhibitor or [S] for stimulatory. (C) High magnification micrographs demonstrate the relative abundance of inflammatory cells in *RNF43* mutant COADREAD tumors, compared to RNF43 WT COADREAD. (D) Violin plots comparing the distribution of leukocyte fraction among LUAD with mutations in *STK11*, *KEAP1*, both (STK11+KEAP1), or neither (WT) in TCGA. The horizontal line indicates the median for each population. Comparisons were made using the Wilcoxon rank-sum test. Significance levels of the associations are labelled as FDR<0.15 (*), FDR<0.05(**), FDR<0.01 (***). (E) Color gradients showing the associations between KEAP1 and STK11 mutation status and leukocyte subtype in TCGA. Values represented are linear regression coefficients. Significance levels of the associations are labelled as FDR<0.15 (*), FDR<0.05(**), FDR<0.01 (***). (F) Overall survival for ICI-treated LUAD patients with *KEAP1* and *STK11* mutations (*KEAP1+STK11*) was significantly worse compared to *KEAP1* alone (log-rank FDR = 0.03), *STK11* alone (log-rank FDR=0.03), or WT (log-rank FDR = 0.01)

In order to assess whether *KEAP1*+*STK11* co-mutation was predictive of ICI outcomes or prognostic of poor outcomes generally, we examined its association with survival outcomes in the TCGA LUAD cohort. This analysis revealed that *KEAP1*+*STK11* status was also associated with worse outcomes in the TCGA LUAD cohort, which was not ICI-treated (HR = 2.5, FDR =0.02, **Supplemental Table 12**), suggesting that *KEAP1*+*STK11* LUAD may be a poor prognosis subgroup, regardless of the treatment strategy.

### IRAGs are associated with cancer-specific features of the immune microenvironment

Immunotherapy outcomes have been associated with features of the tumor immune microenvironment (TME), such as the presence of particular immune cell types^30–34^ and the expression of immune checkpoint molecules^35^. Using mutation data and immune features of the TCGA pan-cancer cohort^36–38^, we examined whether the IRAGs were associated with alterations in the TME (**Methods**).

First, we compared the molecularly-derived leukocyte fraction between IRAG-mutant and WT samples in TCGA, to determine whether IRAG status was associated with overall immune infiltration (**Methods**). Median leukocyte fraction was significantly higher in *NCOR1* (0.24 vs 0.15, p<0.001) and *RNF43* mutant (0.24 vs 0.15, p<0.001) COADREAD samples compared to WT, and significantly lower in *KEAP1* mutant LUAD samples compared to WT (0.26 vs 0.31, p = 0.009) (Figure 3A). KEAP1 mutant LUAD samples also exhibited significantly lower mRNA expression of both stimulatory and inhibitory immune checkpoint molecules compared to WT (e.g. *VTCN1, CD274, CTLA4, CD80, CD86,* all FDR<0.001). Conversely, *RNF43* mutant COADREAD samples had higher mRNA expression of six inhibitory checkpoint molecules, including *VTCN1* (FDR = 0.04), *LAG3* (FDR = 0.05), and *CD274* (FDR = 0.13), features consistent with greater benefit from ICI therapy (Figure 3B, **Supplemental Table 14**). These results also illustrate that IRAGs were associated with different immune features in different cancer types (**Supplemental figures 3A-L**).

We additionally examined hematoxylin and eosin-stained (H & E stain) slides from TCGA (**Methods**) and found evidence of increased lymphocytic infiltration among *RNF43* mutant COADREAD compared to WT (**Figure 5C, Supplemental Table 15**). Qualitative assessment of a selected series revealed greater infiltration of lymphocytes between tumor cells and a higher prevalence of tumor necrosis in *RNF43* mutant cases compared to WT controls, supporting the possibility of RNF43 p.G659Vfs*41 as an immunogenic neoantigen. Most lymphocytes in WT controls were in connective tissue, rather than penetrating the tumor.

Given the poor prognosis associated with *KEAP1*+*STK11* co-mutation, we also analyzed the immune microenvironment of tumors with both mutations (*KEAP1*+*STK11*), either mutation alone (*KEAP1, STK11*), or neither mutation (WT). We found that *KEAP1+STK11* tumors had a significantly lower median leukocyte fraction than WT (0.20 vs 0.31, p <0.001), *STK11* (0.20 vs 0.29, p = 0.01), and *KEAP1* tumors (0.20 vs 0.29, p = 0.04) (Figure 3D). *KEAP1+STK11*, *KEAP1*,and *STK11* tumors were each associated with reduced dendritic cells (p = 0.002, <0.001, <0.001, respectively) and B cells (p = 0.002, <0.001, 0.03, respectively) compared to WT, as determined by gene signature-based deconvoluted cell types (Figure 3E). All three categories of tumors also had broadly decreased immunomodulator expression and immune signature scores (**Supplemental Figure4**).

### IRAGs improve patient stratification beyond TMB and MSI for immunotherapy

Recent retrospective studies utilized TMB to further stratify patients with MSI-H^5^ and microsatellite stable (MSS)^7^ tumors into subpopulations with different immunotherapy outcomes. We sought to enhance patient stratification using combinatorial genomic markers; specifically, we determined whether IRAG mutation status could improve outcome prediction beyond MSI and TMB status (**Methods**).

As expected, the majority of MSI-H tumors were COADREAD samples (n = 29), though there were also a few MSI-H BLCA (n = 8), STAD (n = 7), LUAD (n = 3) and HNSC (n = 2) samples. We stratified the cohort by MSI status (MSI-H or MSS) and conducted multivariable Cox regression analysis (accounting for age, sex, ICI type, and TMB) to determine whether IRAG mutation status was an independent predictor of survival within these subgroups (Figure 4A). Among MSI-H COADREAD, *RNF43* was independently associated with improved survival (HR = 0.11, p = 0.02), while among MSS COADREAD, NCOR1 was independently associated with worse survival (HR = 8.66, p = 0.007) (Figure 4B). Since MSI-H was uncommon in other cancer types, the MSS cohorts were generally similar to the primary cohorts, and all of the other IRAGs remained significantly associated with survival among MSS patients.

**Figure 4.**
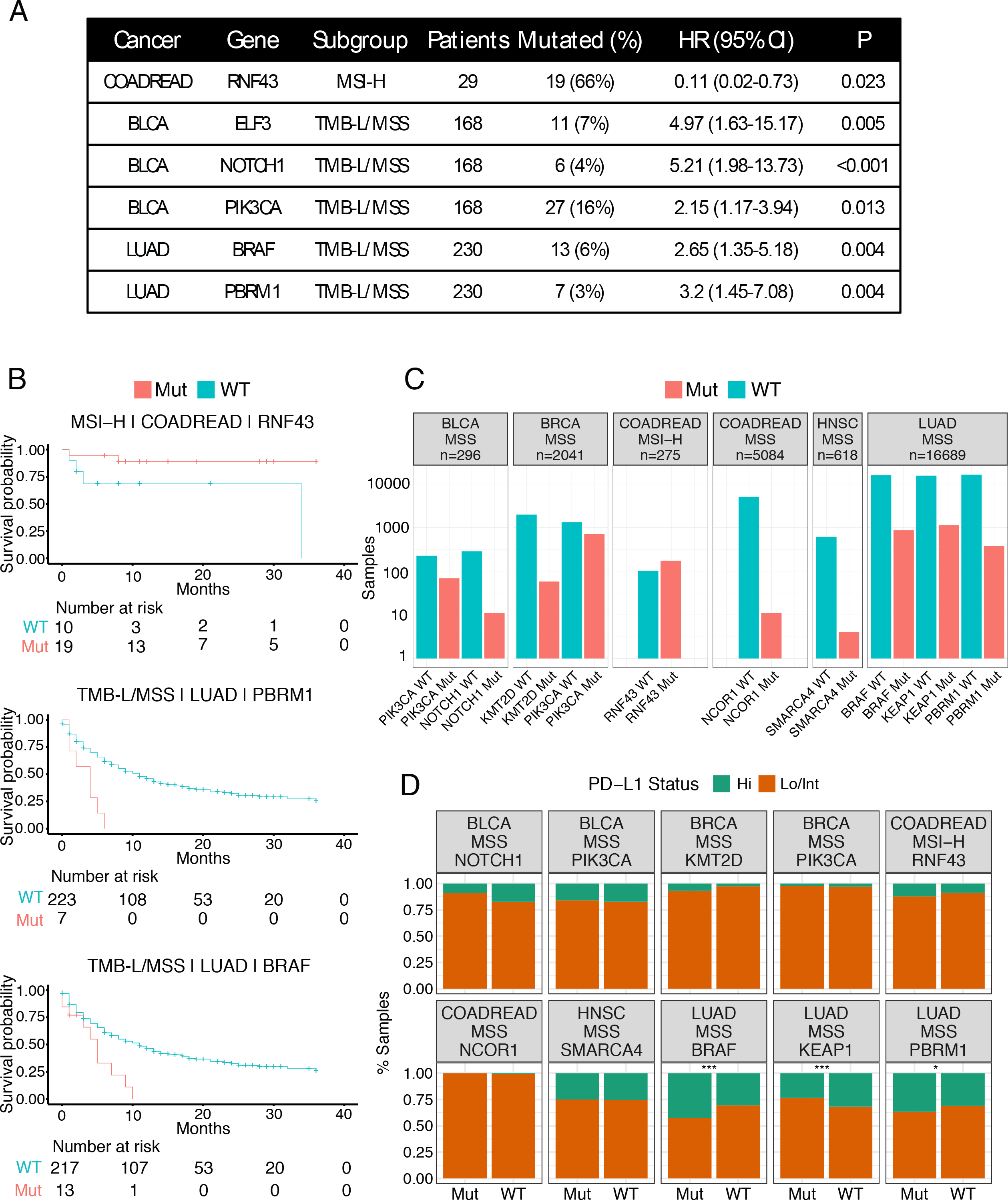
Stratification of immunotherapy response by TMB, MSI and IRAG status. (A) Table showing IRAGs associated (p<0.05) with outcomes in patient subgroups defined by MSI and/or TMB. Hazard ratios (HR) for IRAG mutation status are from multivariable Cox proportional hazards models within the specified patient subgroup. (B) Kaplan-Meier curves illustrating stratification of overall survival using IRAG mutation status within subgroups defined by MSI and/or TMB. (C) The number of IRAG mutations in the Foundation Medicine cohort. (D) The proportion of tumors in Foundation cohort with PD-L1 high or low/intermediate status, by IRAG mutation status. Fisher’s exact test FDR <0.15 (*), <0.05(**), <0.01 (***).

We also investigated whether IRAG mutation status was an informative predictor when stratifying the cohort by both TMB and MSI (Figure 4A, 4B, and **Supplemental Figures 5A-B**). Using the same multivariable Cox regression approach, we identified several independent predictors of survival among MSS/TMB-L tumors: *NOTCH1* (HR = 5.2, p<0.001), *ELF3* (HR = 5.0, p= 0.005) and *PIK3CA* (HR = 2.2, p = 0.01) in BLCA; *BRAF* (HR = 2.7, p = 0.004) and *PBRM1* (HR = 3.2, p = 0.004) in LUAD; *NCOR1* in COADREAD (HR = 8.3, p = 0.02). Among MSS/TMB-H tumors, ROS1 (HR = 3.5, p = 0.008) and *SMARCA4* (HR = 6.9, p = 0.03) in HNSC remained independently associated with survival. Overall, these results demonstrate that IRAGs may provide additional predictive value in stratifying patients who can benefit from immunotherapy.

In order to assess the real-world applicability of the IRAGs, we assessed their prevalence in the Foundation Medicine cohort of 25,003 cases across five cancer types, and examined their association with TMB and PD-L1 expression (as determined by immunohistochemistry). As expected, many IRAGs were mutated in a meaningful fraction of Foundation samples—*RNF43* in 63% of MSI-H COADREAD, *KEAP1* in 6.8% of LUAD—though IRAG mutation prevalence was generally lower in the Foundation cohort than in the Samstein discovery cohort (Figure 4C). PD-L1 expression was increased in *PBRM1* and *BRAF*-mutant LUAD (FDR = 0.09 and <0.001, respectively), while KEAP1 was associated with decreased PD-L1 expression (FDR <0.001), in agreement with immune feature associations that we observed in TCGA (Figures 4C, 3B). KEAP1 in LUAD was also associated with higher median TMB (8.8 vs 7.0 mut/Mb, FDR<0.001),consistent with prior reports.^13^ *RNF43* was not significantly associated with increased PD-L1 expression (FDR = 0.86), but was associated with increased TMB (47.5 vs 41.1 mut/Mb, FDR = 0.8, **Supplemental Table 18, Supplemental Figure 7**) in MSI-H COADREAD.

## Discussion

The majority of cancer patients do not respond to immunotherapy, and our ability to predict clinical benefit is limited. Here we report the identification of 12 ICI-response associated genes (IRAGs) across 6 cancer types, which were predictive of immunotherapy outcomes. The list validates genes(*KEAP1*^12,13^,*PBRM1*^15,16^, *POLE*^17^) and pathways (chromatin remodeling^39^, Wnt signaling^30,40^) that have been previously implicated in ICI benefit in both clinical and preclinical studies, and nominates new candidates for mechanistic functional studies and clinical validation, such as *ELF3, KMT2D,* and the Notch signaling pathway (*NCOR1, NOTCH1*). Notably, we identified *RNF43* as a promising new candidate biomarker for immunotherapy.

To date, genomic predictors of ICI benefit have been analyzed predominantly at the gene level, implying that different variants are collapsed into binary mutant versus wild-type categories. Recent efforts to incorporate specific receptor tyrosine kinase variants into predictive models highlight the potential value of variant-level information^41,42^. Here, we refined our analysis from gene-level to variant-level by considering variant type and clonality as variables that could influence predictive value. Our results demonstrate that the ICI-benefit/resistance phenotype can, in some cases, be ascribed to specific variants, and that variant clonality can be associated with ICI outcomes. Mechanistic studies and larger clinical cohorts will help to elucidate whether variant-level factors improve predictive accuracy versus considering all gene variants equivalently.

The tumor immune microenvironment is a key mediator of ICI outcomes, and we found that IRAG mutations were associated with specific immunologic features, such as leukocyte fraction and immunomodulator expression (Figure 3). *KEAP1* in LUAD was associated with an “immune-cold” microenvironment—with reduced leukocyte fraction and reduced expression of immunomodulators—and worse outcomes. Conversely, *RNF43* in COADREAD was associated with increased leukocyte fraction, increased expression of immunomodulators, and improved survival. These findings lend mechanistic support to the associations between IRAGs and ICI outcomes.

Our results add more detail to prior reports of *KEAP1* and *STK11* associations with worse ICI outcomes^11–13,22,43^. Although we identified *KEAP1* as an IRAG, co-mutation analysis revealed that it was predominantly the *KEAP1+STK11* co-mutated subset that drove the poor outcomes (Figure 3F). We replicated this finding in another ICI cohort (**Supplemental Table 11**), and then tested the association in the non-ICI TCGA cohort (**Supplemental Table 12**), which revealed that *KEAP1+STK11* co-mutated tumors were a poor prognosis subgroup regardless of therapy. Strikingly, *KEAP1* and *KEAP1+STK11* mutated tumors exhibited global downregulation of immunomodulators and low leukocyte fractions (Figure 3B, 3E, 4C) which may underlie the poor prognosis of this tumor subtype, as an immune-infiltrated tumor microenvironment has been implicated in response to both chemotherapy^44^ and immunotherapy^33–35, 45^.

Although some genomic treatment response markers are tissue-agnostic (dMMR/MSI-H, *NTRK* gene fusions), the tissue of origin is often influential via epigenetics and differences in the baseline immune microenvironment. Indeed, we found that IRAGs had tissue-specific associations. *KMT2D, SMARCA4, PIK3CA,* and *PBRM1* were the only IRAGs to be associated with survival in more than one cancer type (Figure 1D). In addition, IRAG mutations were associated with variations in the tumor immune microenvironment across cancer types (**Supplemental Figure 3**). Accounting for tumor-specific effects of mutations may improve immunotherapy outcome prediction further.

In the context of immunotherapy, it has been recently shown that *PBRM1* mutations are associated with better outcomes in KIRC, but only in those previously treated with VEGF inhibitors ^15,16^. We did not identify *PBRM1* as an IRAG in KIRC; however, information on prior therapy was not available for the cohort and was not accounted for. Interestingly, we identified *PBRM1* as a negative predictor of survival in LUAD, which is a novel cancer-specific finding. Larger immunotherapy-treated cohorts are needed to confirm this association.

Notably, four IRAGs are involved in chromatin modification. *PBRM1* and *SMARCA4* are both components of BAF chromatin remodeling complexes, while *KMT2D* is a histone methyltransferase, and *NCOR1* can repress gene expression by recruiting histone deacetylases^46,47^. Epigeneticconsequencesofsomaticmutationsmayinfluenceimmunotherapy responsiveness. A recent study found that Pbrm1-deficient murine melanoma cells were more susceptible to T cell killing and had more accessible chromatin at IFN-γ-responsive sites^39^. Thus, epigenomic analysis of tumors may yield further insight into their biological state and predict immunotherapy responsiveness.

Several IRAGs were predictive of ICI outcomes even after stratifying by TMB and MSI. Among tumors lacking markers of ICI response (i.e., TMB-L/MSS), IRAGs delineated a subset of tumors with even worse outcomes: those with *ELF3, NOTCH1,* or *PIK3CA* mutations in BLCA, and those with *BRAF* or *PBRM1* mutations in LUAD (Figure 4).

Conversely, *RNF43* in COADREAD appears promising as a predictor for improved outcomes. RNF43 is mutated in over 18% of COADREAD and endometrial carcinomas and is strongly linked with MSI-H status in these tumors^28^. Although MSI-H is itself a positive predictor of ICI outcome, *RNF43* mutations were still associated with improved survival among MSI-H COADREAD (Figure 4). *RNF43* encodes an E3 ubiquitin ligase, which negatively regulates Wnt signaling^48,49^. Interestingly,*RNF43*variants have also been proposed as commonly expressed neoantigens for cellular or vaccine-based immunotherapy inCOADREAD^24,25^. RNF43-mutated COADREAD tumors in TCGA had increased leukocyte infiltration and increased expression of immunomodulators, suggesting an “inflamed” tumor microenvironment. Thus, *RNF43* mutations may identify a subtype of MSI-H COADREAD that is poised to respond to ICI treatment.

The reported findings are limited to hypothesis-generating associations, which must be confirmed by prospective and mechanistic studies. This was a sizeable cohort of more than 1,500 ICI-treated patients, but sample size and power were still limited for most cancer types. Overall survival has the advantage of being an objective endpoint, but it can be affected by subsequent treatment, which was not accounted for. Limiting the analyses to the commonly mutated genes reduced the false discovery rate, but may also have omitted less frequently mutated IRAGs (e.g., B2M, JAK2^14^). Certain driver gene mutations which have clinically available treatments (e.g., *ALK, EGFR* in LUAD) may be underrepresented in this cohort, since these patients are unlikely to receive ICI treatment unless targeted therapies have failed. Expanding immunotherapy cohorts with sequencing data will likely reveal additional outcome-associated genes^50^.

In summary, we identified a group of commonly mutated driver genes associated with survival after ICI treatment. Our findings confirmed several previously observed associations and also identified new genes and pathways that may shed light on the response and resistance mechanisms of cancer immunotherapy. Clinically, these IRAGs may also enable more precise stratification of patients for ICI therapy.

## Methods

### Datasets and clinical cohorts

Samstein et al: Clinical and genomic data from the study by Samstein et al. were downloaded from https://www.cbioportal.org/.^21,51,52^ This cohort included 1,661 patients who had received at least one dose of an ICI (targeting PD-1, PD-L1 or CTLA-4) and who had tumor genomic profiling using the commercially available MSK-IMPACT assay, which identifies somatic exonic mutations in a panel of genes. The most recent version of the panel used in the cohort included 468 genes, while previous versions included 341 or 410 genes. Cancer types were defined by aggregating Oncotree codes into TCGA abbreviations. Patients whose cancer type was not known, or with subtypes with 20 or fewer patients in the cohort (e.g. uveal melanoma, chromophobe kidney cancer, papillary kidney cancer), were excluded given inadequate statistical power. Tumor mutation burden for these patients was previously derived by Samstein et al. by normalizing the number of somatic nonsynonymous mutations by the total numberofmegabasessequenced.

Miao et al.: This was a cohort of 249 ICI-treated patients with microsatellite-stable (MSS) solid tumors. Pre-treatment samples were analyzed with whole-exome sequencing (WES). Clinical and genomic data were downloaded fromhttps://www.cbioportal.org/.^15^

Van Allen et al.: This was a cohort of 110 ICI-treated patients with melanoma whose pre-treatment tumors were analyzed using WES. Clinical and genomic data were downloaded from https://www.cbioportal.org/.^19^

TCGA: The Cancer Genome Atlas (TCGA) is a publicly funded project comprising over 11,000 tumors across 33 tumor types, with multiple levels of data including clinical outcomes, sequencing, and expression data (available at https://gdc.cancer.gov/about-data/publications/pancanatlas). A variety of immune features have also been derived from TCGA data^36^, which we combined with mutation data to evaluate the association between somatic mutations and the immune microenvironment. These data were downloaded from https://www.cbioportal.org/.

Foundation Medicine: This cohort comprised 25,003 tumor samples in a commercial cancer sequencing database (Foundation Medicine). Tumor specimens were sequenced using a next-generation sequencing gene panel, and PD-L1 status was determined via immunohistochemistry using Dako 22C3 or Ventana SP142 antibodies. This cohort included five cancer types: COADREAD (MSI-H n = 275, MSS n = 5084), LUAD (n = 16689), BRCA (n = 2041), BLCA (n = 296) and HNSC (n = 618). Cohort level data on IRAG mutation status, PD-L1 status, and median TMB was provided (**Supplemental Table 18**).

### Multivariate Cox regression to identify genes and variants associated with immunotherapy outcome

Cox proportional hazards regression modeling was carried out using the *survival* package in R (R foundation, Vienna, Austria). The primary predictor was each gene’s mutation status; only the top 10% most commonly mutated genes in each cancer type were evaluated, to limit the hypotheses testing to cases with sufficient power. Only nonsynonymous mutations were considered, including both truncating and missense mutations. Covariates were age group at diagnosis, sex, ICI class (anti-CTLA-4, anti-PD-1/PD-L1, or a combination) and TMB. Separate models were constructed for each cancer type. Multiple hypothesis testing correction was performed using the Benjamini-Hochberg procedure to calculate the false discovery rate (FDR) within each cancer type. Thresholds of p<0.05 and FDR<0.2 were used to identify immune ICI-response associated genes (IRAGs). Kaplan-Meier plots were generated using the *survminer* and *ggplot2* packages in R.

A similar analytic pipeline was used to analyze variants for associations with overall survival. The primary predictor was the presence or absence of the specific variant in question (compared to those without the gene mutation, or with other variants in the same gene). Only nonsynonymous variants in the previously identified IRAGs were considered and only variants with more than one occurrence were evaluated. The analysis was stratified by cancer type and IRAGs were tested only in associated cancer types. Covariates were age group at diagnosis, sex, ICI class (anti-CTLA-4, anti-PD-1/PD-L1, or a combination) and TMB. FDR was calculated using the Benjamini-Hochberg procedure within each cancer type. A threshold of FDR<0.2 was used to identify variants associated with ICI outcome. A threshold of p<0.05 was used to identify suggestive associations between variants and ICI outcome.

Cox regression models involving the TCGA, Miao, and Van Allen cohorts were constructed in analogous fashions. Covariates in all models included age, sex and a variable to reflect mutation burden.

### Variant-level analyses

In order to compare the prevalence of specific variants in the IRAG-associated cancer compared to other cancers with the IRAG mutation, we first identified the variants with more than one occurrence in the IRAG-associated cancer. For each variant, we identified all patients who had the IRAG mutation, and constructed a 2×2 table comparing the presence or absence of the variant with the presence or absence of the IRAG-associated cancer type. We then used a one-sided Fisher’s exact test to determine whether the variant was enriched in the IRAG-associated cancer type compared to other cancer types with the same gene mutation.

The variant allele fraction (VAF) was calculated as the number of variant reads divided by the total number of reads at that locus. The tumor purity-adjusted VAF (tpVAF) was obtained by dividing VAF by the histologically-estimated tumor purity. Since there were only 6 copy number changes affecting IRAGs, we reasoned that tpVAF could be used as a proxy for mutation clonality. In cases where a tumor had more than one variant in the same gene, we used the highest tpVAF for that sample. More in-depth methods for estimating the true fraction of cancer cells possessing a variant correct for copy number changes and treat sequencing reads as coming from a binomial distribution.^53^

Treating tpVAF as a continuous variable, we used the *survivalROC* package in R to generate time-dependent ROC curves. We then selected the optimal cutpoint possessing the highest value of Youden’s index.

### Co-occurring and mutually exclusive gene analyses

We used the two-tailed Fisher’s exact test to evaluate for a tendency towards co-occurrence or mutual exclusivity between IRAGs and other mutated genes within each IRAG’s associated cancer type. We performed multiple hypothesis testing correction using the Benjamini-Hochberg procedure to calculate the false discovery rate (FDR) within each cancer type. A threshold of FDR<0.05 was used to identify significant tendencies.

In order to evaluate the effect of co-occurring or mutually exclusive mutations, we first assessed whether these mutations were independently associated with overall survival using a multivariable Cox proportional hazards model adjusting for age at diagnosis, sex, ICI class, and TMB (e.g. OS ~ Mutation + Age + Sex + ICI + TMB), stratified by cancer type. Only those mutations which were independently associated with survival (p<0.05 and FDR<0.2) in this regression were evaluated in the next step (**Supplemental Table 13**). FDR was calculated within each cancer type. Although *RB1* was associated with survival in COADREAD (HR = 66.2, FDR = 0.15), we excluded it from further analyses as there were only 4 *RB1*-mutated COADREAD samples,allofwhichco-occurredwith *NCOR1* mutations.

For each remaining mutation (Gene B), we evaluated whether it confounded the association between the IRAG (Gene A) and overall survival by adding Gene B as an additional covariate to the model (e.g. OS ~ Gene A + Gene B + Age + Sex + ICI + TMB). We considered Gene B a confounder if Gene A was no longer independently associated with survival after Gene B was added to the model.

We also evaluated for significant interactions between IRAGs and the co-occurring genes associated with survival identified above. The Cox proportional hazards model for interaction was constructed as OS ~ A*B + A + B + Age + Sex + ICI + TMB. We used a threshold of p<0.05 to define significant interaction terms.

Validation of the association between *KEAP1+STK11* co-mutation status and worse outcomes was performed in the Miao et al. and TCGA cohorts by classifying LUAD patients into four categories by *KEAP1* and *STK11* mutations status. A multivariable Cox regression model was constructed with *KEAP1/STK11* mutation status as the categorical primary predictor and with wild-type as the reference group. The model was adjusted for age, sex and TMB.

### Determination of gene expression associated with variants

AeQTL was used to perform eQTL analysis on aggregated somatic truncations and missense mutations across 32 cancer types of the TCGA pan-cancer cohort.^37^

For each tested gene region, AeQTL performed a linear regression analysis of RNA-Seq gene expression against genotype and covariates: Gene expression ~ Genotype + Age + Gender + Ethnicity + Cancer subtype + PC1 + PC2 + Young-onset. The linear model was built using the ordinary least squares method with a residual term that follows a normal distribution with a mean of zero and a constant variance.

A separate AeQTL run was performed for each variant type (truncations and missense mutations) in each cancer type, and all the summary files for each variant type were compiled together for multiple hypothesis testing correction using the Benjamini-Hochberg method (FDR<0.05) to generate the final pan-cancer output.

### TCGA somatic mutations and immune features analyses

Somatic mutation calls in the TCGA were obtained from the MC3 project.^38^ Immune features such as leukocyte fraction, immunomodulator expression, gene expression signatures and cell types were derived by Thorsson et al.^36^ We used multivariable linear regression models to test whether mutations in genes of interest were independently associated with immune features, adjusting for age at diagnosis and population substructure (the first two principal components) and stratifying by cancer type. The analysis was conducted using the *glm* function of the base package of the R project (Vienna, Austria). We set the *glm* parameter “family=poisson()” for count data, and “family= gaussian()” for gene expression signatures. The model was:

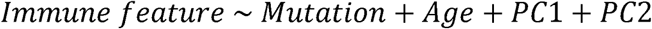

Key immune features were: leukocyte fraction (derived from methylation data), immunomodulator expression (e.g. PD-L1, CTLA-4, etc.), immune expression signatures (e.g. wound healing, TGF-β response, lymphocyte infiltration, macrophage regulation, IFN-γ response), and scores of inflammation, such as the CYT score. Adjustment for multiple comparisons was achieved using the Benjamini & Hochberg method to calculate the false discovery rate (FDR). We used thresholds of FDR<0.15 and FDR<0.05 to identify suggestive and significant associations, respectively, between mutations and immune features.

### TCGA H&E slide review

Whole slide images of hematoxylin and eosin-stained slides from TCGA were available through the Genomic Data Commons Data Portal of the National Cancer Institute (https://portal.gdc.cancer.gov). We selected 10 patients with advanced (T3/T4) *RNF43*-mutated colorectal cancer and 10 patients with advanced *RNF43*-WT colorectal cancers for qualitative comparison. The slides were assessed by a board-certified pathologist using a semi-quantitative scale from 0 to 3, where 0 represented absence of lymphocytic infiltration in the specimen, and 3 represented the highest in the examined series.

### Stratification of outcomes in combination with TMB, MSI and PD-L1 status

MSI high (MSI-H) status was defined as an MSIsensor score ≥ 10.^54^ Samples below this cutoff were considered microsatellite stable (MSS). MSIsensor is an algorithmic method for determining MSI status from paired tumor-normal sequencing data which has been validated in the MSK-IMPACT panel.^54,55^

Tumor mutation burden (TMB) was calculated as the number of nonsynonymous somatic mutations divided by the number of megabases sequenced. TMB thresholds vary. Some studies have used flat TMB cutoffs^7,56^, while others have advocated for cancer-specific thresholds^21^. We used a cutoff of the top 20% of TMB in each cancer type, as Samstein et al. had previously shown this to be effective in this cohort.^21^

To determine the utility of IRAGs in combination with MSI and/or TMB, we tested the IRAG associations with overall survival using multivariable Cox regression in cancer-specific cohorts stratified by MSI and TMB. Cohorts with fewer than 30 patients were excluded. The primary predictor was the IRAG, while other covariates were age, sex, ICI type and TMB (e.g. OS ~ IRAG + Age + Sex + ICI + TMB).

### PepQuery peptide search

We used PepQery, a peptide-centric search algorithm that enables identification of target peptides within proteomic data using a novel DNA or protein sequence of interest.^26^ The PepQuery online portal is available at http://pepquery2.pepquery.org/ (accessed 3/25/2020). In order to identify evidence of the *RNF43* p.G659Vfs*41 variant protein, we searched PepQuery using “Prospective_Colon_VU_Proteome” as the MS/MS dataset, “Protein Sequence” as the target event, “PQRKRRGVPPSPPLALGPRMQLCTQLARFFPITPPVWHILGPQRHTP” as the input protein sequence, “hg38_RefSeq_20190910” as the reference database, and “HyperScore” as the scoring algorithm, with unrestricted modification filtering enabled. We cross-referenced patient IDs from the spectrum titles with genomic data from the CPTAC cohort (available at http://linkedomics.org/cptac-colon/, accessed 3/25/2020) to identify patients with the variant in question.

### Statistical analysis

Proportions of categorical variables were compared using the chi-squared test or Fisher’s exact test. The chi-squared test was used for contingency tables larger than 2×2. One-sided Fisher’s exact tests were used when testing for the enrichment and two-sided tests were used when testing for either enrichment or depletion.

Unless otherwise stated, multiple hypothesis testing correction was achieved using the Benjamini-Hochberg method. Unless otherwise stated, we used thresholds of FDR<0.2 and p<0.05 to determine significance. Tests with FDR<0.2 but p>0.05 were not considered significant.

### Patient Consent (Foundation Medicine)

Approval for this study, including a waiver of informed consent and a HIPAA waiver of authorization was obtained from the Western Institutional Review Board (Protocol No. 20152817). Consented data that can be released is included in the paper and its supplementary files. Patients were not consented for release of raw sequencing data.

### Genomic method (Foundation Medicine)

Genomic data were collected as part of routine clinical care using a targeted comprehensive genomic profiling assay in a Clinical Laboratory Improvement Amendments (CLIA)-certified, College of American Pathologists (CAP)-accredited, New York State-approved laboratory (FoundationOne^®^, Cambridge, MA, USA), as previously described^57^.

### PD-L1 staining (Foundation Medicine)

PD-L1 staining was performed at Foundation Medicine (Cambridge, MA, USA). PD-L1 status was determined through immunohistochemistry performed on formalin-fixed paraffin-embedded (FFPE) tissue sections, with the use of the commercially available antibody clones 22C3 (Dako/Agilent, Santa Clara, CA, USA) or SP142 (Ventana, Tuscon, AZ, USA), as previously described^6^.

## ACKNOWLEDGEMENTS

The authors wish to acknowledge those involved in the generation and curation of data supplied to the public by The Cancer Genome Atlas, the Clinical Proteomic Tumor Analysis Consortium, Memorial Sloan Kettering Cancer Center, and Dana Farber Cancer Institute. Most importantly, we thank the patients who have generously contributed their data to these various projects.

## DATA AND SOFTWARE AVAILABILITY

Datasets are available online, as described in the Methods. R code for the analyses can be found at https://github.com/Huang-lab/IRAG-2020

## COMPETING FINANCIAL INTERESTS

The authors declare no competing financial interests. L.P. has received honoraria unrelated to the topic of this paper from Astra Zeneca, Merck, Novartis, Bristol-Myers Squibb Genentech, Eisai, Pieris, Immunomedics, Seattle Genetics, Clovis, Syndax, H3Bio and Daiichi. LAA, GMF and JFH are employees of Foundation Medicine Inc and stock holders of Roche Holdings AG.

## CONTRIBUTIONS

TJ and KH conceived and designed the research and analyses. TJ, TQ, GD, JFH, and LAA conducted the analyses. MS and CC conducted the pathology review. TJ and KH interpreted the results and drafted the manuscript. KH supervised the study. All authors read, edited, and approved the manuscript.

